# The Aging Brain Matrix: Cross-Species Evidence Linking *FREM3* to Brain and Cognitive Aging

**DOI:** 10.64898/2026.10.27.762265

**Authors:** Kevan P. Clifford, Thomas D. Prevot, Akiko Sumitomo, Srija Das, Yashika Bansal, Michael Marcotte, Runhao Zhou, Jonas Yeung, Megan Ding, Daniel Felsky, Brian J. Nieman, Mounira Banasr, Etienne Sibille, Toshifumi Tomoda, Yuliya S. Nikolova

**Author notes:** **Corresponding authors:** Toshifumi Tomoda, MD & PhD CAMH, 250 College Street Toronto, ON, M5T1R8, Canada, Yuliya Nikolova, PhD CAMH, 250 College Street Toronto, ON, M5T1R8, Canada. Equal contributions as senior authors (Toshifumi Tomoda, Yuliya S. Nikolova).

## Abstract

The human extracellular-matrix-related *FREM3* gene is strongly age-dependent, enriched in age-sensitive excitatory long-range-projecting neurons, and implicated in neurodegenerative diseases. To establish the first translationally relevant causal evidence for the gene’s roles in brain structure and cognition, we developed an excitatory-neuron-specific *Frem3* conditional knockout mouse model (*Frem3*-cKO) and computed genetically regulated expression of *FREM3* (*FREM3*- GReX) in humans, enabling convergent neurostructural and cognitive characterizations. *Frem3*- cKO mice showed widespread lower global and regional brain volume in middle age vs young adulthood, consistent with drastic age-dependent volume loss absent in wildtype mice. These effects were strongest in cognitively relevant cortical regions and paralleled by lower cognitive performance. In humans, lower *FREM3-*GReX was associated with lower total surface area, which further mediated an indirect link to lower cognitive performance across multiple age-sensitive domains. We provide the first translationally relevant cross-species evidence supporting a key role for *FREM3/Frem3* in maintaining brain and cognitive health across aging.

## 1. Introduction

Understanding the drivers of brain aging is key to identifying novel targets to prevent and treat age-related cognitive decline and neurodegeneration. The brain matrisome, consisting of the Extracellular Matrix (ECM) and ECM-affiliated proteins, plays a key role in the maintenance of brain tissue structural integrity (Dityatev et al., 2010) and is almost universally impaired in neurodegenerative disease (Sun et al., 2021). Identifying specific matrisome proteins, whose age-related change is causally linked to structural and cognitive changes, is a crucial first step to developing novel matrisome-based interventions to prevent age-related neurocognitive decline.

We recently conducted the largest transcriptome-wide study of cortical age-dependent gene expression across the normal adult lifespan in a non-clinical postmortem cohort of 214 individuals aged 20-90, with balanced sampling across ages. The *FREM3* (Fras1-related Extracellular Matrix Protein 3) gene, emerged as the top matrisome gene most downregulated with age across the two prefrontal cortical regions sampled (Brodmann areas 47 and 11) (C.-W. Lin et al., 2021). Similar age-dependent reductions in *FREM3* expression were reported in independent, smaller postmortem brain tissue datasets (GTEx Consortium, 2013; Hu et al., 2018). Further, we previously demonstrated that *FREM*3 alleles associated with lower *FREM3* mRNA levels in young adults, effectively shifting *FREM3* expression towards an older-age-like state, are associated with blunted corticolimbic system reactivity and slower processing speed (Y. Nikolova et al., 2015)— a neurobehavioural profile more commonly observed in late life. Together, these studies suggest that lower *FREM3* mRNA levels are among the most distinguishing features of an aging brain matrisome and may drive “accelerated” or premature brain aging in younger individuals.

Consistent with links between accelerated brain aging and neurodegenerative disease (Cao et al., 2010; Glorioso et al., 2019; Peng et al., 2021), reduced *FREM3* mRNA has also been found in postmortem corticolimbic brain tissue of individuals with Alzheimer’s disease (AD), relative to age-matched cognitively intact controls (Agora, 2026a; Hokama et al., 2014). In line with its extracellular-matrix-related function and predicted role in cell adhesion (National Library of Medicine, 2026), *FREM3* is classified under the ‘Structural Stabilization’ biological domain within the TREAT-AD framework that maps genes to AD-relevant biological processes (Cary et al., 2024). Notably, among genes assigned to this domain, *FREM3* consistently ranks among the six most downregulated in frontal and temporal cortical regions in AD (Agora, 2026b; Britton et al., 2025). Beyond AD, reduced *FREM3* has been identified as part of a pan-neurodegenerative transcriptomic cortical response shared between Parkinson’s and Huntington’s disease (Labadorf et al., 2017). Conversely, higher levels of *FREM3* mRNA have been associated with neuroprotective or “brain reserve” phenotypes such as larger overall brain size independent of chronological age (Fenu et al., 2018; Roe et al., 2011; Seidlitz et al., 2023). Together, these studies suggest that lower *FREM3* levels may be associated with risk for neurodegeneration, while higher *FREM3* levels may contribute to neuroprotection and brain reserve. Yet, strikingly, causal evidence for the gene’s involvement in these phenotypes is lacking.

In the adult human brain, *FREM3* mRNA follows a primarily corticolimbic expression pattern with the strongest expression levels in prefrontal and frontopolar regions (Berg et al., 2020; The Human Protein Atlas, 2026) and is enriched in age-sensitive long-range-projecting excitatory neuronal subtypes (Hodge et al., 2019) associated with cognition (Berg et al., 2020; Galakhova et al., 2022) and cognitive resilience to aging (Consens et al., 2022). Consistent with a potential structural support role, FREM3 consists primarily of Chondroitin Sulfate Proteoglycans (CSPG) repeats (Pavlakis et al., 2011; Blum et al., 2025; InterPro entry IPR039005), thought to mediate cell adhesion, including cell-cell and cell-matrix interactions (Avram et al., 2014).

The *Frem3* gene is the only homolog of the human *FREM3* gene in mice, showing 69% identity and 80% homology on the protein level, and is similarly expressed in excitatory long-range projecting neurons (Tasic et al., 2016). This cross-species conservation supports the use of mouse models to investigate the neurobiological consequences of altered *Frem3* expression. Prior constitutional *Frem3* genetic knockout (KO) studies, however, have not identified any major abnormalities (Kiyozumi et al., 2020). This is broadly consistent with several established brain- matrisome KO models, whose deficits are only revealed through more sensitive neurostructural and behavioural assays (Brakebusch et al., 2002; Montag-Sallaz & Montag, 2003; Weber et al., 1999; Zhou et al., 2001). Thus, targeted assays sensitive to the specific neurostructural and cognitive phenotypes most likely to arise from *Frem3* loss may be necessary to reveal its functional consequences.

*FREM3*’s selective enrichment in long-range-projecting excitatory neurons and its association with overall brain size (Seidlitz et al., 2023) suggest that *FREM3* may contribute to neurostructural properties spanning multiple spatial scales. Mesoscale, whole-brain approaches such as small-animal MRI may therefore be particularly well-positioned to capture its effects, from regional structural alterations to broader spatial patterns of brain-wide structural variation. Importantly, the conservation of *FREM3/Frem3* expression across the human and mouse brain, together with the broad translational correspondence of MRI-derived phenotypes across species (Y. Nikolova et al., 2018), provides a powerful framework for linking experimentally defined effects of *Frem3* in mice to individual differences in human brain structure and aging.

Thus, to study the effects of *Frem3* in a translationally relevant way, we developed a first-of-its-kind conditional *Frem3* knockout model (*Frem3*-cKO), where *Frem3* was removed specifically from corticolimbic excitatory neurons to model human expression patterns. Here, we introduce this novel model, its molecular validation, as well as its neurostructural and cognitive effects, measured via whole-brain structural MRI (sMRI) and behavioral analysis, respectively, in young adult and late middle-aged mice. To enable reverse translation across species, we leveraged the late middle-aged UK Biobank cohort to compute genetically regulated expression of *FREM3* and tested its links to brain structure and cognitive performance. We hypothesized that in both mice and humans, lower *Frem3/FREM3* expression would be associated with older-age-like phenotypes, including smaller global and regional brain structural measures, alongside worse cognitive outcomes. Together, these complementary approaches provide, to our knowledge, the first cross-species investigation into the role of the age-sensitive *Frem3/FREM3* gene in shaping aging-related brain structure and cognitive outcomes.

## 3. Results

### 3.1 Mouse Results

#### 3.1.1 Development of a novel corticolimbic CaMKII^+^-neuron-specific *Frem3* conditional KO (*Frem3*-cKO) mouse model

To achieve cell type-specific manipulation of *Frem3* gene expression in mice, we used CRISPR/Cas9 to design a conditional allele of *Frem3* using a pair of loxP sites to flank the entire mature coding region spanning immediately downstream of the signal peptide (SP) within the exon-1 through the 3’-untranslated region in the exon-8, as detailed in Methods (**Figure 1A**). The resulting ‘floxed’ locus has the 16-amino acid insertion at the end of the SP. We first validated that this genetically-modified SP could function as efficiently as the parental SP, using the myc-tagged recombinant Frem3 proteins (either harboring the wild-type SP of 27 amino acids or the modified SP of 43 amino acids (=27+16)) expressed in the heterologous system. Surface biotinylation of the transfected cells followed by Western blot analysis confirmed that the equivalent fraction of the expressed, biotinylated Frem3 proteins migrated at the expected size of an apparent molecular mass of ∼31.1 kDa (**Figure 1B**), suggesting that the modified SP did not affect translocation of the recombinant proteins across the plasma membrane and that removal of the modified SP occurred efficiently upon translocation to the endoplasmic reticulum. Next, a test cross of mice with a floxed allele (*Frem3*^flox/+^) with a germline Cre line of mice led to generation of a null allele (*Frem3*^null/+^), demonstrating successful integration of loxP sites in a *cis* configuration at the *Frem3* locus.

**Figure 1.**
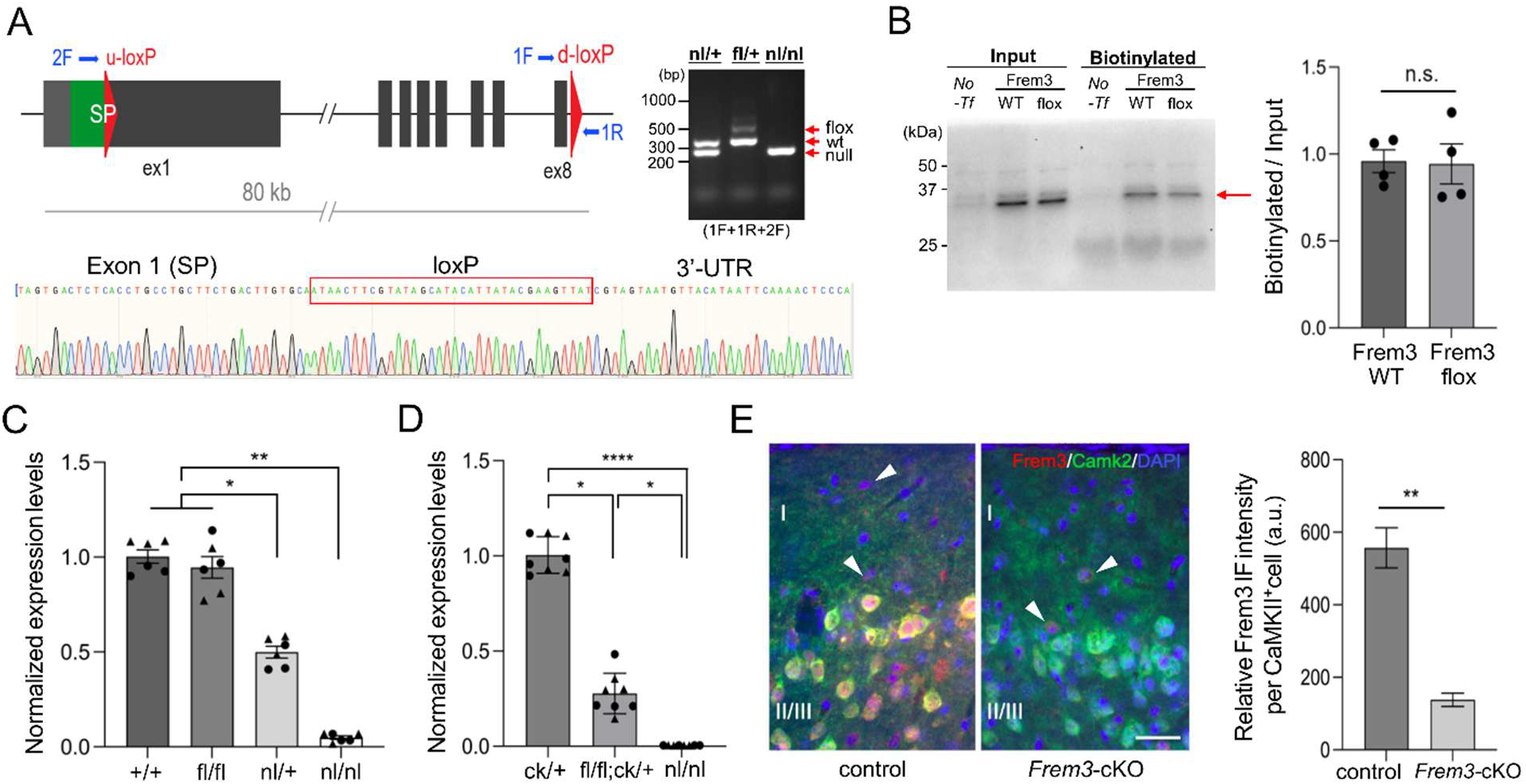
Development of Frem3-cKO mice. (A) Schematic illustration of CRISPR/Cas9 strategy for designing the conditional *Frem3* knockout allele on mouse chromosome 8. The upstream and downstream loxP sites (u-loxP, d-loxP) are inserted in exon-1 and after exon-8, respectively. PCR primers used for genotyping are denoted as 1F, 1R and 2F, differentiating the wild-type (318-bp), floxed (477-bp) and null alleles (232-bp), as shown in the gel image. Sanger sequencing confirmed generation of a null allele upon germline Cre-mediated recombination of the floxed allele. (B) Surface biotinylation followed by avidin beads purification and Western blot, done in quadruplicate, confirmed equivalent membrane surface localization of the recombinant Frem3 protein with the wild-type SP (27 amino acids) or the genetically-modified SP (43 amino acids), as assessed by Mann-Whitney U test (n.s. = not significant). (C) qPCR confirmed no significant change in *Frem3* mRNA expression in PFC of homozygous floxed (*Frem3*^fl/fl^) mice, and an expected reduction in heterozygous (*Frem3*^nl/+^) (by ∼50%) or homozygous null (*Frem3*^nl/nl^) mice (∼100%) as compared with wild-type mice (*Frem3*^+/+^). N=6 (⚫=3♂; ▴=3♀)/group (3 months old). (D) qPCR showed ∼73% reduction in *Frem3* mRNA in PFC of *Frem3*-cKO (*Frem3*^flox/flox^;Camk2a^Cre/+^ (= fl/fl;ck/+)) as compared with control mice (*Frem3*^+/+^;Camk2a^Cre/+^ (= ck/+)). N=8 (⚫=4♂; ▴=4♀)/group (3 months old). *p<0.05; **p<0.01; ****p<0.0001 (Kruskal- Wallis with Dunn’s post-hoc test). (E) Immunofluorescence analysis showed marked reduction in Frem3 protein level (red) in CaMKII^+^ pyramidal neurons (green) in layer II/III of PFC in *Frem3-* cKO as compared with Camk2a^Cre/+^ control mice (3 months old). Bar: 50 μm.

To assess the impact of loxP insertion on *Frem3* gene expression, we produced homozygous floxed (*Frem3*^flox/flox^) or null (*Frem3*^null/null^) mice via intercrossing of respective heterozygous mice and quantitated *Frem3* mRNA expression in prefrontal cortex (PFC) as compared with that in wild-type control mice (*Frem3*^+/+^). qPCR assay showed that *Frem3* expression in *Frem3*^flox/flox^ mice was equivalent to that in wild-type mice and that *Frem3*^null/+^ or *Frem3*^null/null^ mice had a half or virtually none *Frem3* expression, respectively (**Figure 1C**), together demonstrating that the loxP insertion had no discernible effect on *Frem3* gene expression and that the floxed allele had high fidelity of Cre-mediated recombineering capacity.

Next, we generated corticolimbic excitatory neuron-specific KO mice (*Frem3*^flox/flox^;Camk2a^Cre/+^), hereinafter referred to as *Frem3*-cKO, via crossbreeding with Camk2a-Cre mice (*i.e.*, ♂*Frem3*^flox/+^ x ♀*Frem3*^flox/+^;Camk2a^Cre/+^). qPCR assays showed that *Frem3*-cKO exhibited a ∼73% reduction in *Frem3* mRNA level in PFC compared to the littermate control (*Frem3*^+/+^;Camk2a^Cre/+^) (**Figure 1D**), in line with predominant *Frem3* expression in cortical excitatory projection neurons detected in single-cell RNA-seq (Tasic et al., 2016). Consistently, immunofluorescence assays showed that Frem3 protein expression was predominantly detected in CaMKII^+^ neurons in the layer II/III of PFC in control mice, whereas the Frem3 immunoreactivity was selectively lost in CaMKII^+^ neurons of *Frem3*-cKO mice while weaker levels of Frem3 expression remained detectable in other cell types (arrowhead) for both genotypes (**Figure 1E**). These results support pyramidal neurons as the primary cell type of *Frem3* expression in the adult mouse cortex.

#### 3.1.2 Mouse neurostructural effects

We observed a highly significant age-related reduction in total brain volume for *Frem3*-cKO mice (late middle-aged < young adults; t = −3.274; p = 0.006, controlling for sex). In comparison, control mice did not show a significant age-related change in total brain volume (t = −0.852, p = 0.41; interaction t = −1.862; p = 0.0736). Post hoc regional analyses indicated widespread significantly lower volume for 98 of 180 regions in late middle-aged vs young adult *Frem3*-cKO mice (all pFDR < 0.05), while no regions showed significant age-related differences in control mice after FDR correction (all pFDR ≥ 0.371).

The 98 regions showing selective age-dependence in *Frem3*-cKO mice revealed pervasive lower volume across the isocortex, encompassing 42 of 44 isocortical ROIs, with strongest effects in frontal association and medial orbital cortices. Further isocortical reductions extended across all primary and secondary motor, somatosensory, and visual cortices, the insular area, as well as the temporal and posterior parietal association areas. Beyond the isocortex, significantly lower volume was observed in 14 of 16 cerebellar gray matter subfields, all parahippocampal and 5 of 20 hippocampal formation subfields, and select subcortical regions including the thalamus, amygdala, claustrum, striatum, and hypothalamus. In addition to gray matter effects, 4 of 21 white matter and 6 of 23 cerebellar white matter subregions demonstrated significantly lower volume. Conversely, brainstem, pituitary, and deep cerebellar regions showed no significant volumetric differences, alongside select olfactory, basal ganglia, and most hippocampal formation subregions. Significant regional *Frem3*-cKO effects are illustrated in **Figure 3A**. Full statistical results for regional within-genotype brain volume effects are available in Supplementary Data 1.

Notably, however, only 24 of the 98 age-sensitive regions in *Frem3*-cKO showed nominal age-by-genotype interaction effects, mainly spanning isocortical regions (frontal association, medial orbital, and motor cortices, as well as select cingulate subregions) and 12 of 16 cerebellar gray matter and three cerebellar white matter regions. Regions demonstrating both significant lower within-*Frem3*-cKO volume and nominal age-by-genotype interactions are illustrated in **Figure 2B** and detailed in **Table 1**. Full statistical results for all age-by-genotype brain volume interactions are available in **Supplementary Data 1**.

**Figure 2.**
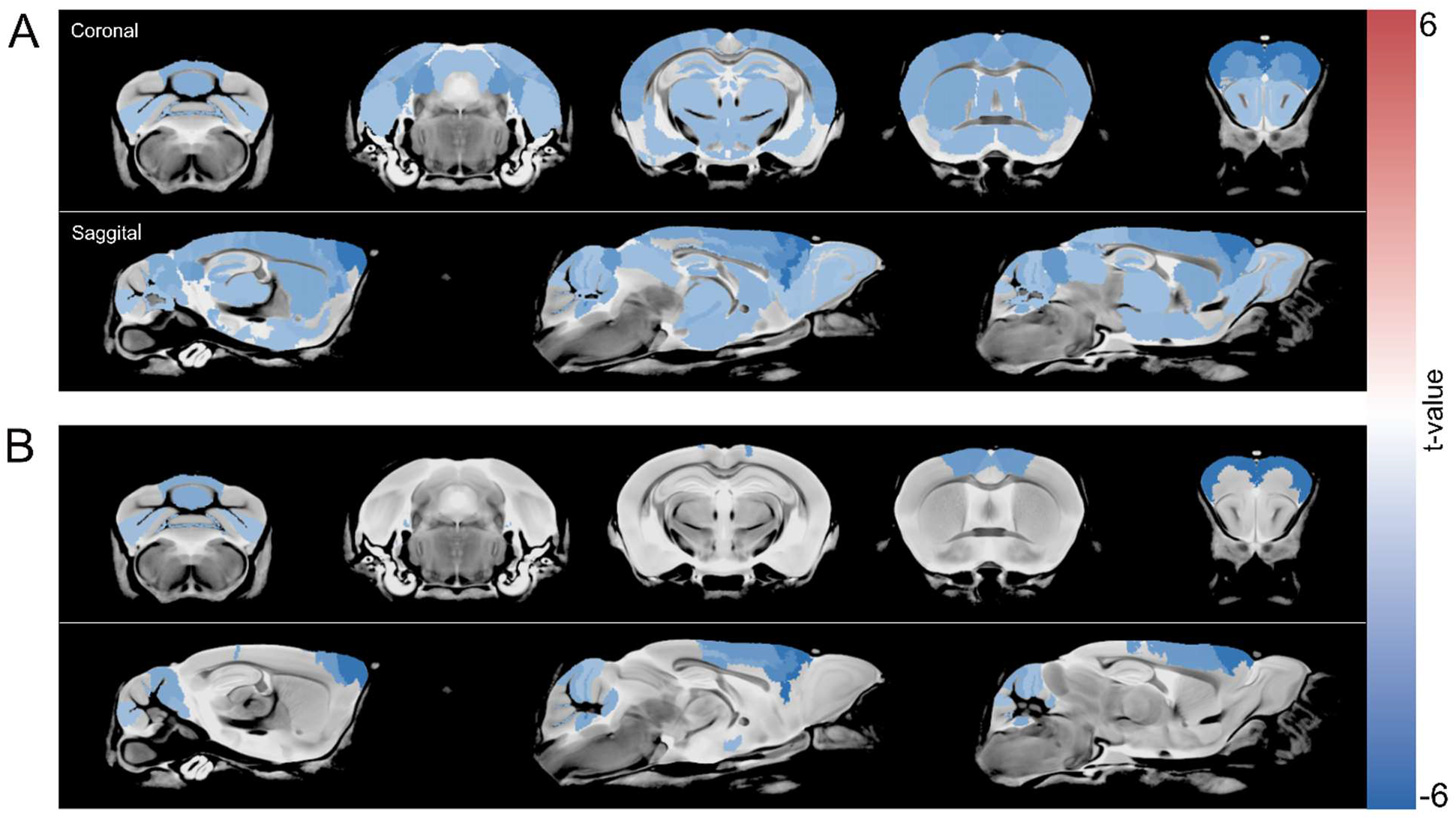
Significant aging-related lower brain volume within Frem3-cKO mice and overlapping age-by-genotype effects. Panel A depicts regions with significant brain volume loss between young adult and late middle-aged *Frem3*-cKO mice (n = 98 regions; all pFDR < 0.05) on a t-value scale. Of these regions, 24 showed additional age-by-genotype interaction effects (B) (p < 0.05).

**Table 1.** Within-*Frem3*-cKO regions showing both significant age-related lower volume and corresponding nominal age-by-genotype effects.

| Region | Within<br>*Frem3*-cKO<br>t-val | Within<br>*Frem3*-cKO<br>pFDR | Age × genotype<br>interaction p-val |
| --- | --- | --- | --- |
| **Isocortex** |  |  |  |
| Frontal association cortex | -5.394 | <0.001 | 0.007 |
| Medial orbital cortex | -5.569 | <0.001 | 0.029 |
| Cingulate cortex area 24a.1 | -3.893 | 0.021 | 0.005 |
| Cingulate cortex area 24b | -3.559 | 0.021 | 0.033 |
| Cingulate cortex area 32 | -4.592 | 0.021 | 0.020 |
| Primary motor cortex | -3.950 | 0.021 | 0.028 |
| Secondary motor cortex | -4.047 | 0.021 | 0.038 |
| Parietal cortex posterior area rostral part | -3.530 | 0.021 | 0.046 |
| Cingulate cortex area 24b.1 | -3.113 | 0.027 | 0.034 |
| **Cerebellum** |  |  |  |
| Lobule 10 nodulus | -4.946 | <0.001 | 0.003 |
| Lobules 4 5 culmen ventral and dorsal | -3.414 | 0.021 | 0.002 |
| Lobule 6 declive | -3.337 | 0.021 | 0.034 |
| Simple lobule lobule 6 | -3.399 | 0.021 | 0.002 |
| Lobules 1 2 lingula and central lobule ventral | -3.253 | 0.023 | 0.020 |
| Lobule 9 uvula | -3.282 | 0.023 | 0.016 |
| Flocculus FL | -3.148 | 0.027 | 0.008 |
| Lobule 8 pyramis | -3.021 | 0.030 | 0.023 |
| Anterior lobule lobules 4 5 | -2.966 | 0.031 | 0.027 |
| Crus 1 ansiform lobule lobule 6 | -2.882 | 0.035 | 0.019 |
| Copula pyramis lobule 8 | -2.788 | 0.039 | 0.012 |
| Paramedian lobule lobule 7 | -2.601 | 0.045 | 0.031 |
| **Cerebellar white matter** |  |  |  |
| Lobule 10 white matter | -3.442 | 0.021 | 0.005 |
| Lobules 4 5 white matter | -2.563 | 0.048 | 0.012 |
| **Hypothalamus** |  |  |  |
| Medial preoptic nucleus | -2.910 | 0.033 | 0.041 |

#### 3.1.3 Mouse Y-maze alternation task

In contrast to these neurostructural effects, which were largely age-dependent, cognitive impairment was consistent across both age groups. Specifically, a two-way mixed-effects model showed a significant main effect for genotype (*F*(1, 30) = 38.13, p < 0.0001), with no age (F(1, 23) = 7.82 × 10⁻⁵, p = 0.9930) or age-by-genotype interaction (*F*(1, 23) = 1.184, p = 0.2877) effects. Šídák-corrected post hoc comparisons showed that *Frem3*-cKO mice had significantly lower alternation rate in the Y-maze, suggestive of a robust working memory impairment, at both 4 months (p < 0.0001) and 12 months (p < 0.0001) compared to control mice (**Figure 3**).

**Figure 3.**
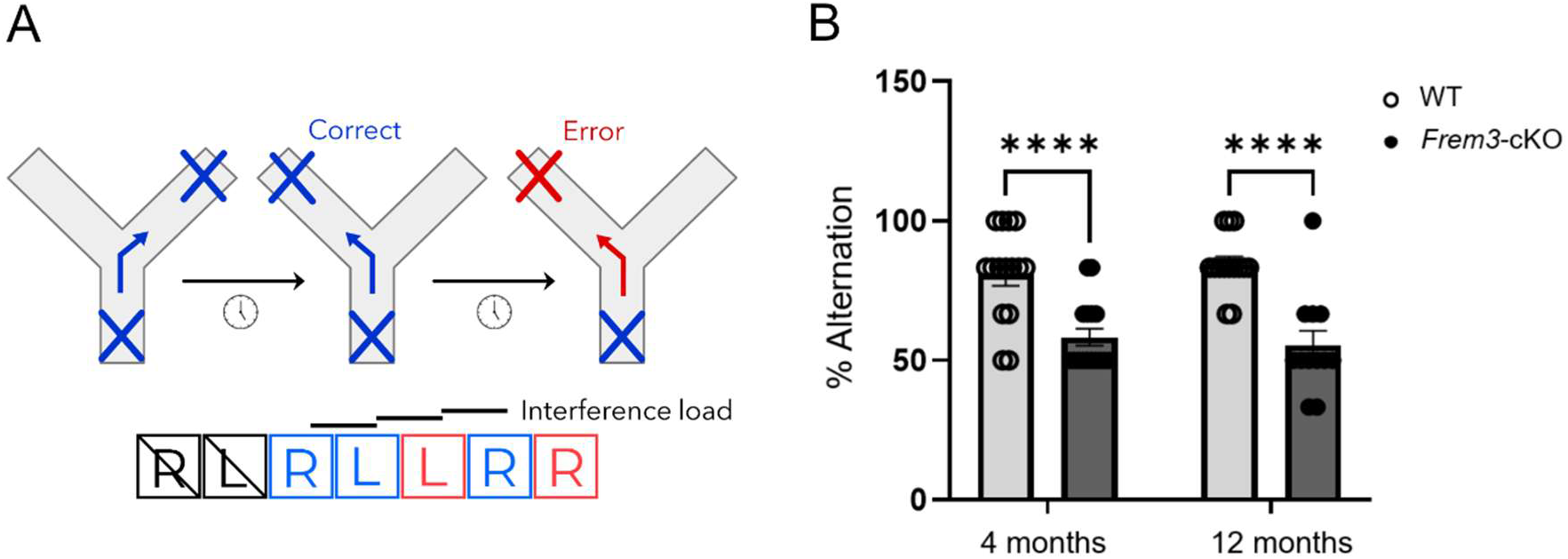
Y-maze alternation task performance in Frem3-cKO mice. In the full sample (B), Šídák-corrected post hoc comparisons showed *Frem3*-cKO mice to have significantly lower alternation rates than control mice at both 4 months (p < 0.0001) and 12 months (p < 0.0001). 50% represents chance level.

### 3.2 UK Biobank Results

#### 3.2.1 Human Brain Structure and *FREM3*-GReX Associations

In our final sample of 31,384 (16,392 females, aged 46-82; mean = 64.1, ±7.65), lower *FREM3*- GReX was strongly associated with lower total surface area (SA) (t = −3.118; pFDR = 0.006). In contrast, total cortical volume (CV) and mean cortical thickness (CT) did not show a significant association (both pFDR = 0.34). No significant *FREM3*-GReX-by-age interactions were observed across global brain structure measures (pFDR range: 0.858–0.917) suggesting the effect is consistent across the mid-to-late life age range represented in this cohort.

Consistent with the global effect on SA, widespread associations between lower *FREM3*-GReX and lower regional SA were also observed. The strongest effects were localized to the occipital cortex, with markedly lower SA in the bilateral cuneus and pericalcarine cortices, alongside the left lingual cortex (t range = −3.679 to −5.382; all pFDR < 0.0001). Additional FDR-significant effects occurred in the left lateral occipital, right lingual, and right isthmus cingulate (all pFDR < 0.03). Similar FDR-trend-level associations were present in the left pars triangularis, precentral, and isthmus cingulate regions, as well as the right lateral occipital (all pFDR < 0.08). Regional *FREM3*-GReX and SA associations are depicted in **Figure 4**. No significant *FREM3*-GReX-by-age interactions were observed for SA across regions (all pFDR ≥ 0.97).

**Figure 4.**
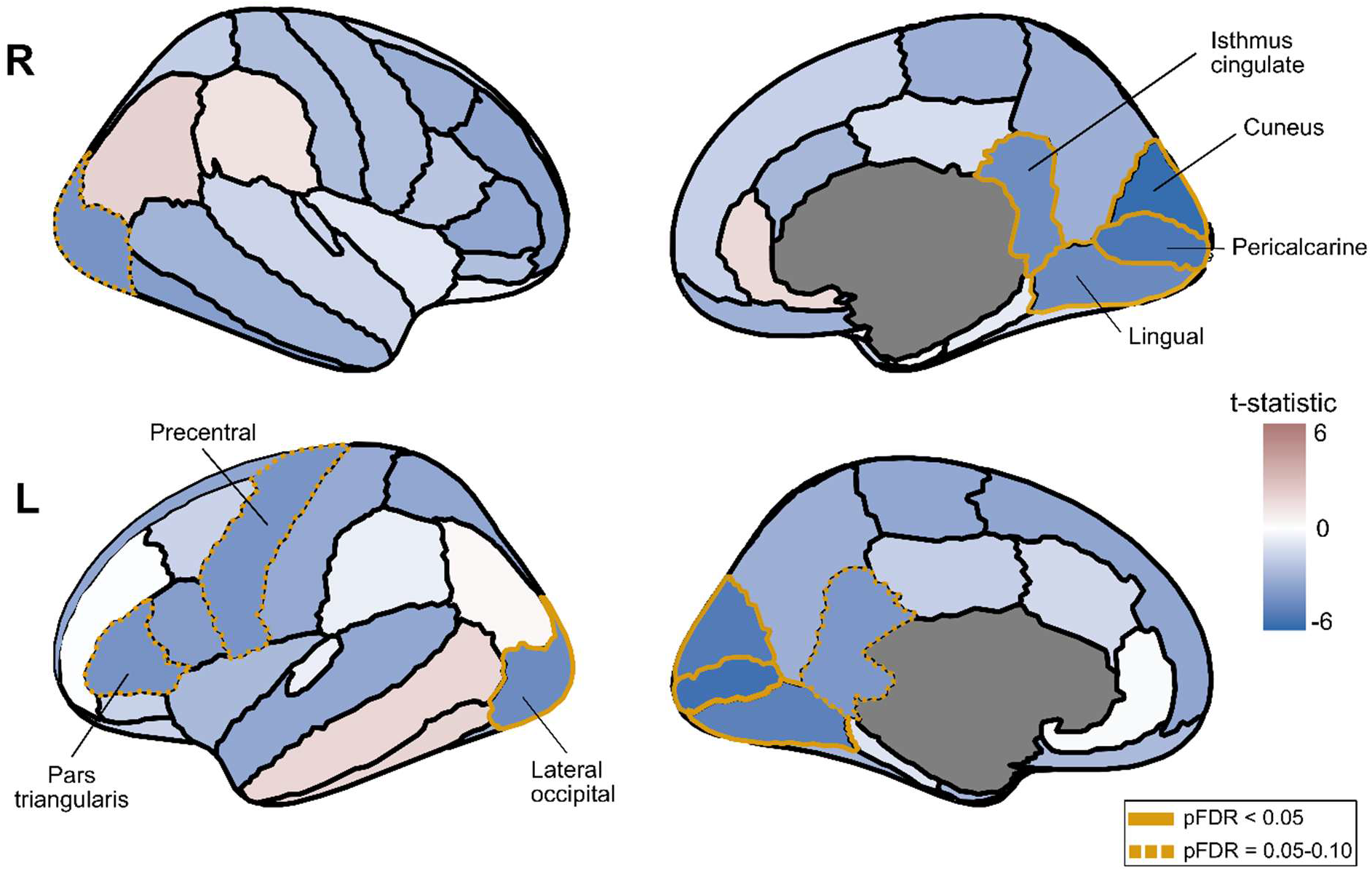
Associations between cortical surface area and FREM3-GReX. Associations between lower genetically regulated *FREM3* expression and regional surface area are depicted across right (R) and left (L) hemispheres of the Desikan-Killiany-Tourville atlas. Blue indicates negative associations (lower surface area) while red indicates positive associations (greater surface area) on a *t*-value scale. Thick orange outlines indicate statistically significant regions (pFDR < 0.05) while thin orange outlines indicate trend-level effects (pFDR = 0.05–0.10).

Exploratory analyses of *FREM3*-GReX and cortical volume effects motivated as a translational comparison to our mouse brain volume measures, revealed concordant spatial and directional patterns with the SA findings including FDR-significant effects in five of the same ROIs (bilateral cuneus and pericalcarine cortices, and right isthmus cingulate (t range = −2.895 to −4.728; all pFDR < 0.05). In contrast, exploratory effects on CT were weak, directionally inconsistent, and did not survive FDR correction. Full statistical results for all *FREM3*-GReX and neurostructural associations are available in **Supplementary Data 2**.

#### 3.2.2 Human Cognitive Performance and *FREM3*-GReX Associations

Direct associations between lower *FREM3*-GReX and cognition were directionally consistent with worse performance but showed no statistically significant effects. The strongest effect was a trend-level association for an executive function measure (Tower rearrange, t = −1.75; p = 0.08), while effects for the other tasks were weaker (p ≥ 0.13). Similarly, no significant *FREM3*-GReX-by-age interactions for cognitive performance were observed (pFDR range: 0.53–0.822).

Critically, however, mediation analyses showed significant indirect effects of *FREM3*-GReX on cognitive performance via total cortical surface area for all cognitive tasks except for reaction time. For the Tower Rearranging Task, a significant indirect effect occurred (ACME=−0.00190, 95% CI [−0.00372, −0.00052], p=0.002), as depicted in Figure 5. Similar significant indirect effects occurred across 7 of the 8 cognitive tasks (all pFDR < 0.05), with the exception of Reaction Time (pFDR = 0.692). Indirect mediation of *FREM3*-GReX effects on cognitive performance through total SA are illustrated in Figure 5A-B. Full statistical results for all *FREM3-*GReX and cognitive performance analyses are available in **Supplementary Data 3**.

**Figure 5.**
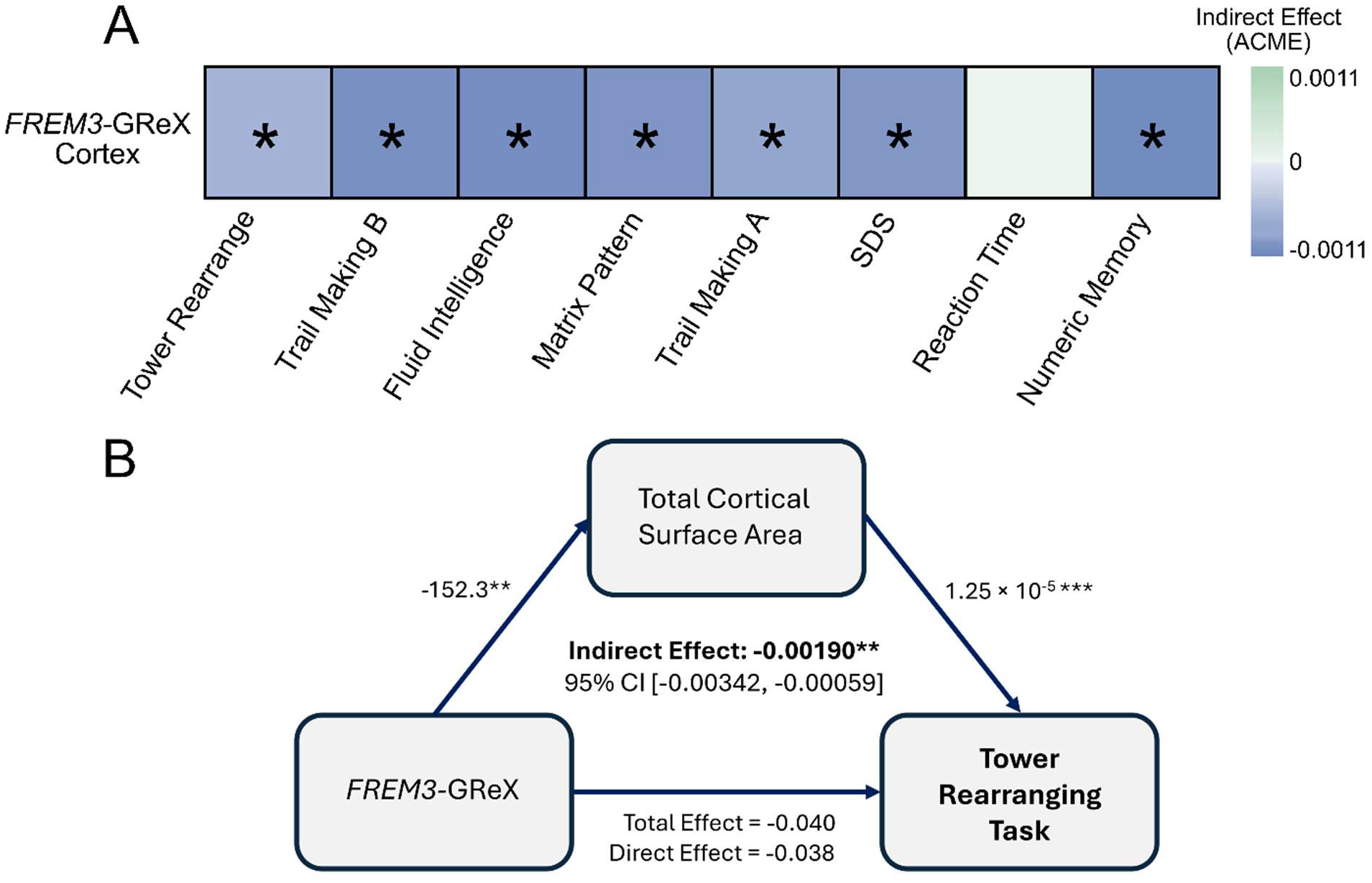
Associations between cognitive performance and FREM3-GReX. (A) Blue indicates worse cognitive performance associated with lower genetically regulated *FREM3* expression while green indicates better performance, mediated by lower total surface area, shown on an estimated ACME color scale (top right). (B) Representative figure depicting the significant mediation effect, which emerged for the Tower Rearranging Task. Similar significant indirect effects were observed for all cognitive tasks after FDR correction, except for Reaction Time (full statistical details in **Supplementary Data 3**). ACME = Average Causal Mediation Effect.

## 4. Discussion

Here, we show convergent neurostructural and cognitive changes associated with lower corticolimbic excitatory neuronal *Frem3/FREM3* expression using a novel *Frem3*-cKO mouse model in parallel with human *FREM3*-GReX associations. *Frem3*-cKO mice showed significant age-related losses in total brain volume and widespread isocortical volume reduction, most strongly in age-sensitive frontal and orbital regions, alongside significant impairment in a working memory task in young and middle-aged animals. In humans, consistent with mouse findings of global brain volume reduction, lower *FREM3*-GReX was also associated with lower total surface area, driven by marked bilateral effects across occipital and isthmus cingulate regions. Furthermore, lower *FREM3*-GReX was consistently indirectly associated with lower cognitive performance via reduced total cortical surface area. Together, these findings provide the first causal evidence linking lower *Frem3* expression to age-related neurostructural and cognitive deficits, while supporting converging correlational effects in humans.

*Frem3*-cKO mice showed markedly lower global brain volume in middle-aged vs young adult animals relative to control mice, which showed no age-related difference. The age-related volumetric differences in *Frem3*-cKO mice was striking on a regional level, with more than half of all ROIs showing lower volume after FDR correction. Among the 24 regions with strongest age-by-genotype effects, most striking volume loss occurred in the frontal association and medial orbital isocortical regions. These frontal regions may be translationally relevant to human *FREM3-* enriched regions integral to long-range cortico-cortical integration underlying higher-order cognition (Haber et al., 2022; Klein-Flügge et al., 2022). Steeper age-related decline was also observed in the motor cortices, including the medial precentral cortex, a key hub for frontal-motor integration that is implicated in age-related motor and executive dysfunction (Barthas & Kwan, 2017; Inoue & Nishimune, 2023). Notably, volume loss also occurred in regions not targeted by the corticolimbic knockout, suggesting *Frem3* may produce distal effects consistent with a role in long-range connectivity. For instance, in contrast to our previous work showing cerebellar preservation across healthy mouse aging (Clifford et al., 2023), much of the cerebellum in *Frem3*- cKO mice displayed age-related volume differences. Overall, these findings highlight distributed cortico-cerebellar circuits spanning motor and frontal association cortices as particularly vulnerable to steeper age-related volumetric decline following loss of *Frem3* expression in corticolimbic excitatory neurons.

An additional 74 regions showed FDR-significant age-related volume loss in *Frem3*-cKO mice without corresponding FDR-significant age effects in control mice and without age-by-genotype effects. Most notably, midlife volume reductions in *Frem3*-cKO mice encompassed nearly the entire isocortex (42 of 44 regions), translationally aligning with human *FREM3*’s cortex-wide expression patterns and its role in higher-order sensory and cognitive processes (Berg et al., 2020). These additional isocortical regions included all cingulate subdivisions, the insular area, and the adjacent claustrum and striatum (implicating cortico-basal ganglia pathways) (Gehrlach et al., 2020), and all sensory cortices. Notably, the observed hippocampal and parahippocampal loss deviates from patterns of healthy aging preservation observed in our previous work and others (Clifford et al., 2023; Maheswaran et al., 2009).

In parallel to these neurostructural effects, young adult and late middle-aged *Frem3*-cKO mice from an independent cohort demonstrated significantly impaired working memory performance compared to controls, suggesting a considerable role for *Frem3* in supporting working memory function across adulthood. Notably, behavioral deficits were present in young adulthood and did not reveal subsequent progressive impairment in middle age. This may reflect a floor effect in the Y-maze task, as both young adult and late middle-aged *Frem3-*cKO mice performed near the 50% chance level, limiting sensitivity to detect worsening performance (Prevot et al., 2021). Future studies should leverage more sensitive paradigms to explore potential progressive mid-to-late life memory deficits.

In humans, neurostructural effects showed partial convergence with mouse findings. Lower *FREM3*-GReX was most significantly associated with lower total SA in mid-to-late life, consistent with *FREM3*’s role in global structural support. Furthermore, lower total SA mediated an indirect link between lower *FREM3*-GReX and lower performance across all cognitive tasks except reaction time. This suggests that *FREM3*-GReX may influence overall cognitive performance more strongly through its associated effects on global brain structure.

Regionally lower SA was associated with lower *FREM3*-GReX most prominently in bilateral occipital regions. This encompassed the primary visual cortex (pericalcarine), the cuneus, implicated in visual attention-shifting and visual working memory (Simpson et al., 2011), the lingual gyrus, supporting higher-order visual processing and visual memory encoding (Palejwala et al., 2021), and the lateral occipital cortex, which supports shape-based object representation (Grill-Spector et al., 2001). Together, these observed effects span primary and higher-order visual regions that support visuospatial representations and their integration into higher-order association networks (Bettcher et al., 2016; Cox et al., 2019; Jung & Haier, 2007).

Typical occipital SA trajectories show gradual aging-related decline (Long et al., 2012; Nyberg et al., 2023), with some evidence for non-linear, steeper aging-related reduction (Storsve et al., 2014). In this context, the marked lower SA observed across occipital regions may reflect aspects of aging-related pathology. In particular, these patterns align with reports of lower SA in case-control studies and predictive models of major depressive disorder (MDD) (Belov et al., 2024; Couvy-Duchesne et al., 2018; Harris et al., 2022; Matsumoto et al., 2023), a neuropsychiatric disorder characterized by premature brain aging (Han et al., 2021; Koutsouleris et al., 2014). This may broadly align with previous transcriptomic studies associating altered *FREM3* expression with MDD in postmortem corticolimbic brain tissue (Chang et al., 2014; Gryglewski et al., 2022). Interestingly, findings of a large-scale ENIGMA-MDD cortical meta-analysis closely mirrored these spatial patterns, including lower total surface area driven by bilateral lingual and cuneus effects, alongside left lateral occipital and precentral regions, comprising an adolescent/young adult MDD signature (Schmaal et al., 2020). In this context, this regional overlap may reflect a shared set of cortical regions vulnerable to accelerated aging-like change, consistent with MDD’s broader associations with premature brain aging.

Beyond occipital effects, the isthmus cingulate cortex (ICC) showed bilateral lower SA associations. As a posterior default mode network (DMN) region, the ICC links the posterior cingulate to the parahippocampal gyrus, supporting internally directed cognition and episodic memory (Kollenburg et al., 2025). Notably, lower SA in the ICC has been linked to Alzheimer’s disease in case-control and Mendelian randomization studies (C. Li et al., 2011; Wu et al., 2021; Yang et al., 2019), as well as conversion from mild cognitive impairment to Alzheimer’s disease (W. Lin et al., 2020; Ren et al., 2024). Furthermore, lower ICC functional activity is shown to contribute to altered DMN patterns characteristic of Alzheimer’s disease (G. Li et al., 2025; Zhu et al., 2013). Taken together, *FREM3*’s neurostructural influence in this region may contribute to its existing transcriptomic associations with case-control Alzheimer’s disease (Britton et al., 2025; Hokama et al., 2014).

While *Frem3*-cKO and lower predicted *FREM3*-GReX in humans produced generally converging neurostructural phenotypes, the nature of reduced *Frem3/FREM3* expression in these constructs is important to contextualize further. Our knockout mouse model afforded loss of function of *Frem3* in virtually all CaMKIIα^+^ cells generally representing the excitatory neurons in cortical and limbic regions, resulting in overall disruption of *Frem3* expression by ∼73%. In contrast, while the *FREM3*-GReX model performed strongly, it ultimately captures only the cis-genetically regulated component of expression (∼15%) and cannot capture non-genetic regulation (age, environment, disease). Further, since to our knowledge no excitatory-neuron-specific *FREM3* eQTLs have been identified, our *FREM3*-GReX estimate is based on bulk cortical tissue, limiting its cell-type specificity. This highlights constraints of correlational in vivo human imaging-genetics and the complementary value of experimental mouse models. Despite these limitations, it is striking that significant associations do occur between GReX of a single gene and a complex phenotype such as SA, particularly when indexing naturally occurring variability in gene expression. This, coupled with *Frem3*-cKO data, suggests that *FREM3* likely plays an important role in shaping cortical morphology in late middle age.

Our cross-species comparison highlights both shared and divergent neurostructural patterns of lower *Frem3*/*FREM3* expression. Notably, both models suggest a role for *Frem3*/*FREM3* in supporting global brain structure through lower total volume/SA in late midlife. Regional concordance included marked occipital effects in late middle-aged humans mirrored by significant volume loss across all visual cortices in *Frem3*-cKO mice. Additionally, bilateral human ICC effects corresponded to volume loss across all cingulate subregions in *Frem3*-cKO. More broadly, occipital and ICC regions are embedded in long-range association circuits, suggesting *Frem3/FREM3* may preferentially support regions underlying integrative connectivity (Mitchell et al., 2018; Palejwala et al., 2021). Overall, these converging effects strengthen evidence for *Frem3*/*FREM3*’s biological influence in these regions, and prioritize them for future molecular follow-up. Conversely, partially diverging patterns occurred in frontal regions, wherein humans showed widespread but largely non-significant lower SA compared to pronounced volume loss in *Frem3*-cKO. Given the modest variance captured by *FREM3*-GReX, a construct with stronger expression perturbation may have yielded starker human frontal SA effects.

This study is not without limitations. First, we compared differences between young adult and late middle-aged *Frem3*-cKO mice, whereas future work should incorporate old-age and maturational comparisons to fully characterize lifespan effects of *Frem3*-cKO. Secondly, behavioral testing and ex vivo neuroimaging were conducted in independent mouse cohorts, not allowing for within-subject structural and behavioral analyses. Therefore, the findings are interpreted as convergent group-level effects, and subtle cohort-specific differences cannot be fully discounted. In parallel, because *FREM3*-GReX only captures the genetically regulated component of expression (∼15%) and lacks cell-type specificity, it is considerably less sensitive to human associations than stronger causal effects in *Frem3*-cKO mice. Furthermore, given the cross-sectional nature of the UK Biobank, the observed *FREM3*-GReX effects on SA could also reflect additional of developmental influence of lower genetically regulated *FREM3* expression, warranting future investigation in developmental cohorts. Lastly, UK Biobank cognitive measures are brief and self-administered, likely reducing sensitivity to cognitive effects. Future work should prioritize comprehensive, well-validated cognitive assessments.

Limitations aside, this study provides the first causal evidence for age-dependent effects of reduced *Frem3* expression on brain structure and working memory in mice, alongside converging correlational evidence in humans. This cross-species neurostructural mapping of *FREM3*-linked effects will inform future work in characterizing targetable *Frem3/FREM3*-related molecular pathways for treatment and prevention of age-related neurocognitive disorders.

## 2. Methods

### 2.1 Mouse Study

#### 2.1.1 Development and validation of a novel *Frem3* conditional knockout model

*Frem3* conditional knockout mice (*Frem3*-cKO) were generated on a C57BL/6 background using the Cre/LoxP site-specific recombination system coupled with CRISPR/Cas9 methodology, in collaboration with The Centre for Phenogenomics (Toronto, ON). Murine *Frem3* gene is comprised of 8 exons, with the exon-1 containing the start codon, 27-amino acid long of the signal peptide (SP) for the plasma membrane surface translocation, as well as the majority (>90%) of the *Frem3* coding sequence, while the coding exons 2 through 7 all contain triplet numbers of nucleotides and the final exon-8 has a stop codon. Due to this unusual genomic organization, we designed a conditional allele of *Frem3* using a pair of loxP sites to flank the entire mature coding region spanning immediately downstream of the SP within the exon-1 through the 3’-untranslated region in the exon-8 (**Figure 1A**). Accordingly, the upstream guide RNA (gRNA-U: 5’- GCTTCTGACTTGTGCACTGA -3’ on the sense strand of mouse chromosome 8) was designed to insert a loxP site within the first exon of *Frem3* (between the amino acid 27 and 28), in a way that the inserted loxP sequence (34-bp) plus a 14-bp stuffer sequence (5’- CCGGTGGCTCCGGT -3’) can be translated to yield the 16-amino acid sequence (ITSYSIHYTKLSGGSG) and fused in-frame immediately downstream of the SP (amino acids 1–27) and also in-frame with the subsequent coding sequence of the exon 1 (amino acids 28 onwards). The glycine/serine flexible amino acid stretch at the end of this 16-amino acid insertion would mitigate possible steric hindrance to ensure full functionality of the modified SP and proper protein translocation across the plasma membrane. The downstream guide RNA (gRNA-D3: 5’- TATGTAACATTACTACGGTA -3’ on the antisense strand of mouse chromosome 8) was designed to insert a second loxP site 581-bp downstream of the *Frem3* stop codon within the final exon. To ensure that a pair of loxP sites were successfully inserted in a *cis* configuration on the same chromosomal strand, mice with a floxed allele (*Frem3*^flox/+^) were bred with a germline Cre line of mice (E2a-Cre, Jackson Lab stock# 003724) to confirm generation of a null allele (*Frem3*^null/+^). Intercrossing heterozygous null mice produced homozygous null mice with an expected Mendelian ratio of ∼25%, which were confirmed viable up to 22 months of age with no overt anatomical or physical deficits, consistent with a previous report on the generation of another *Frem3* null mice by 5-bp deletion from *Frem3* exon-1 (Kiyozumi et al., 2020). Wild-type, floxed and null alleles were differentiated via genomic polymerase chain reaction (PCR) using the following primers (1F, 1R, 2F), which gave 318-bp (1F & 1R) from the wild-type allele, 477-bp (1F & 1R) from the floxed allele, and 232-bp (2F & 1R) from the null allele, respectively, and the identity of each allele were verified by Sanger sequencing (**Figure 1A**).

1F: Forward primer after Exon-8: 5’- CAC AAT TGA TGA GTG CCC TAA CTG -3’
1R: Reverse primer after Exon-8: 5’- GAC AGC CAG AAC TAA GTA GAA GGA C -3’
2F: Forward primer in Exon-1: 5’- GGC ATC CTC TAG ATA TAT CAA CTC -3’

The resulting floxed allele was further validated at the transcript level via the reverse transcription-quantitative PCR (qPCR), using SYBR Green supermix (Bio-Rad) and the following primers spanning the exons 5 and 6 (amplicon size: 111-bp).

F(ex5): 5’- ACG GTC ATC TGC TCT ACT CG -3’
R(ex6): 5’- GTG GAG CAT GCT GGT ATT ATC -3’

To validate the functionality of the genetically-modified SP with the extra 16-amino acid insertion (corresponding to the upstream loxP and the stuffer sequences), murine *Frem3* cDNAs (amino acids 1–300) tagged with a myc-epitope at the carboxy termini, with or without the 16-amino acid insertion, were subcloned in the pRK5 expression vector and transfected in HEK 293T cells, which were then analyzed by surface biotinylation to confirm plasma membrane surface translocation of the recombinant proteins, essentially as described (Sumitomo et al., 2018). In brief, cells were incubated with ice-cold PBS containing EZ-Link Sulfo-NHS-SS-Biotin (Pierce^TM^) for 30 min with gentle rocking at 4°C, then lysed and incubated with NeutrAvidin beads. After extensive washing, proteins adsorbed were eluted from the beads in loading buffer containing dithiothreitol, heated for 5 min at 95°C and then analyzed by Western blot using a myc antibody (mouse monoclonal [9E10], 1 µg/ml, Calbiochem).

To develop experimental cohorts for the corticolimbic excitatory neuron-specific conditional KO of *Frem3* (*Frem3*-cKO), male *Frem3* floxed mice were crossed with female breeders harboring Camk2a-Cre (Jackson Lab stock# 005359), which express Cre recombinase specifically in CaMKIIα^+^ forebrain excitatory pyramidal neurons, avoiding germline recombination of the floxed allele and hence non-specific KO in peripheral tissues (Luo et al., 2020). Levels of Frem3 protein expression per CaMKII^+^ cell were evaluated by indirect immunofluorescence using anti-FREM3 antibody (1:100, rabbit polyclonal, Sigma-Aldrich) and anti-CaMKII antibody (1:50, mouse monoclonal, 6G9, Novus) and analyzed by ImageJ.

#### 2.1.2 Animal Groups

Young adult (3–4 months old) and late middle-aged (12–16 months old) mice comprised excitatory neuron-specific *Frem3*-cKO mice (*Frem3*^flox/flox^;Camk2a^Cre/+^) (n = 16) and littermate control (*Frem3*^+/+^;Camk2a^Cre/+^) (n = 16) mice, balanced by sex. Mice were housed under a 12/12-hour light/dark cycle (7am – 7pm light phase) at consistent temperature (20–23°C) with access to food and water. Imaging and behavioural analyses were conducted in separate cohorts matched for age. Given the ex vivo design, imaging analyses were conducted using a between-subject framework. Animal use and testing procedures were conducted in accordance with the Canadian Council for Animal Care with approval from the institutional animal care committee.

#### 2.1.3 Ex vivo MRI-derived brain volume

Brain volume analyses were conducted in a separate cohort of 16 *Frem3*-cKO mice and 16 littermate control mice. For each genotype, animals were evenly distributed across sex and age, with eight young adult (3-4 months old) and eight late middle-aged (13-16 months old) mice. Mice were anesthetized and perfused with a paraformaldehyde solution (4%) containing a Gadolinium contrast agent (ProHance 2mM). Ex-vivo MRI was performed on intact brains using a 7.0 Tesla MRI scanner (Bruker Ltd.) at the SickKids Mouse Imaging Centre (MICe) facility, as described in our previous work (Y. S. Nikolova et al., 2018). We acquired a T2-weighted, 3-D fast spin-echo sequence, with a cylindrical acquisition of k-space, (TR=350ms, TEs=12ms per echo for 6 echoes, FOV= 20 x 20 x 25 mm^3^, matrix size = 504 x 504 x 630), with 40 µm isotropic voxel size (Spencer Noakes et al., 2017). Images were nonlinearly registered to the DSURQE atlas (Chakravarty et al., 2012; Dorr et al., 2008), and brain volumes were extracted from 185 regions. For all MRI analyses, ROI volumes were averaged across hemispheres in order to minimize multiple testing burden.

#### 2.1.4 Y-maze Alternation Task

Working memory was tested using the Y-maze Alternation Task, as in Prevot et al. (2018, 2021). The apparatus consisted of a three-armed black PVC maze. Each arm was 26cm long, 8cm wide, and 13cm high with a sliding door, separated by 120°. To habituate the mice to the maze, 10 minute free-exploration sessions were conducted over two consecutive days. On the third day, mice were placed in a designated start-arm and underwent a training session consisting of seven consecutive trials, whereby mice alternated between the two goal arms with an inter-trial interval (ITI) of 30 seconds. 24 hours later, the same training task was performed but with the ITI increased to 60 seconds to increase task difficulty. To dissociate working memory deficits from potential loss of motivation, an additional 8th trial was conducted after the 7^th^ trial with a shorter ITI (5 seconds). Animals that failed to alternate in the 8th trial were excluded. Working memory performance was indexed as the percentage of alternations out of 6 scored trials, where a statistically significant drop toward ∼50% accuracy is considered a deficit. The Y-maze test is repeatable, which allowed testing in the same young and middle-aged mice. Mice excluded for failing the 8^th^ trial included 1 female control at 4 months, and at 12 months, 2 male controls, 1 male *Frem3*-cKO, and 3 female *Frem3*-cKO mice.

#### 2.1.5 Statistical Analyses – Mice

##### 2.1.5.1 Mouse Molecular Analyses

Levels of surface biotinylation of the recombinant Frem3 proteins and immunofluorescence data were analyzed by Mann-Whitney U-test. Gene expression levels via qPCR were analyzed by Kruskal-Wallis with Dunn’s post-hoc test.

##### 2.1.5.2 Mouse Brain Volume Analyses

Given the ex vivo design, imaging analyses were conducted using a between-subject statistical framework. First, we tested whether global brain structure showed differential age-related change using a linear model with an age-by-genotype interaction for total brain volume in the full sample, with sex as a covariate. Subsequently, we evaluated within-genotype total and regional brain volume differences using linear models, with sex as a covariate. A false discovery rate (FDR) of 5% was applied to regional analyses (180 regions (ventricles excluded)) to account for multiple comparisons. Since interactions typically require over four times more power to detect than main effects (Cohen, 1977; Galea et al., 2020) and up to 16 times larger study samples (Durand, 2013) these were tested post hoc and results considered exploratory.

##### 2.1.5.3 Mouse Y-maze Alternation Analyses

Working memory performance on the Y-maze alternation task was analyzed in the combined male and female samples of Frem3-cKO and control mice at ages 4 and 12 months of age. A two-way mixed-effects ANOVA was modelled with age, genotype, and age-by-genotype as fixed-effects and subject as a random effect and subsequently corrected using Šídák’s multiple comparisons test. To examine potential sex-specific effects, the same analysis was then performed separately within male and female samples.

### 2.2 UK Biobank Study

#### 2.2.1 UK Biobank Study Population

The UK Biobank is a large population-representative cohort of ∼500,000 participants recruited in the United Kingdom between 2006-2010, with extensive genotypic and phenotypic data, including multimodal neuroimaging and cognitive testing (Sudlow et al., 2015). In the present study, we leveraged genome-wide data, as well as T1-weighted structural MRI-derived phenotypes and cognitive performance measures available during the imaging visit. Genetically regulated expression (GReX) of *FREM3* was computed in 425,630 participants, as described below. After removing related individuals with kinship coefficients > 0.2 and retaining only those with complete neuroimaging data, the final sample size comprised 31,384 mid-to-late life participants (aged 46- 82; mean = 64.1, ±7.65; 16,392 females).

#### 2.2.2 Genetic data and Quality Control (QC)

DNA was extracted from blood samples and whole-genome genotyping was performed using the UK BiLEVE and the UK Biobank Axiom arrays. Genotypes were imputed by the UK Biobank using the HRC and UK10K reference panels. Full details of UK Biobank genetic processing and procedures are described elsewhere (Bycroft et al., 2018). Analyses were restricted to participants that passed UK Biobank QC, which imposed a missing genotype rate of 0.1, Hardy-Weinberg equilibrium of 1e-15, mach-r2-filter imputation quality of 0.8, and a minor allele frequency of 0.01, all within a European population. In addition to UK Biobank QC, we excluded variants with one or more character allele codes, and merged individual chromosomes using PLINK v1.94 (Purcell et al., 2007). Lastly, we accounted for genetic relatedness by excluding individuals with a kinship coefficient > 0.2 while retaining one individual per related pair.

#### 2.2.3 Genetically Regulated Gene Expression

Genetically regulated expression of *FREM3* (*FREM3*-GReX) was computed using PrediXcan (Gamazon et al., 2015). Given *FREM3*’s widespread cortical expression patterns, we leveraged reference cis-eQTL weights from GTEx v8 cortex eQTL models (GTEx Consortium, 2013). An elastic net regression model identified a subset of 22 cis-eQTL SNPs most predictive of previously measured *FREM3* expression. The SNPs were matched by rsID in QC’d UK Biobank genotype data (22/22 available SNPs used) and extracted from QC’d UK Biobank PLINK genotypes (GRCh37) which were used as input for individual-level *FREM3*-GReX estimates. The cortex *FREM3* model demonstrated strong predictive performance in cross-validation (pred_perf_R^2^ = 0.1498; p = 9.65 × 10⁻⁸), explaining approximately 15% of variation in previously measured *FREM3* expression in postmortem tissue. *FREM3*-GReX values were subsequently z-score normalized using the scale() function in R.

#### 2.2.4 MRI Acquisition and Image-derived Phenotypes

The UK Biobank brain MRI protocol was performed using a 3 Tesla Siemens Skyra scanner with VD13 software and a 32-channel head coil, implemented on identical equipment and software across three sites. T1-weighted images were acquired using an MPRAGE sequence at 1mm isotropic resolution. Complete scanner protocol details can be found at http://biobank.ctsu.ox.ac.uk/crystal/refer.cgi?id=2367. UK Biobank image acquisition and processing pipelines are described in detail elsewhere (Alfaro-Almagro et al., 2018; Miller et al., 2016). For our purposes, we utilized T1-weighted images where estimated cortical gray matter was used to automatically generate “imaging-derived phenotypes” (IDPs) values for cortical surface area of 62 regions of the Desikan Killiany-Tourville atlas parcellation (Alexander et al., 2019). Additionally, cortical volume (as a translational comparison to mouse brain volume) and cortical thickness were used in exploratory analyses. Global measures were derived from regional IDPs by summing cortical surface area and volume across regions, and computing mean cortical thickness across regions.

#### 2.2.5 Cognitive Assessments

Cognitive performance tasks were collected during the neuroimaging visit (instance 2). We assessed associations between lower *FREM3*-GReX and performance across age-sensitive cognitive domains, including executive function (Trail Making Test B, n = 21,257; Tower Rearranging Test, n = 20,866), fluid intelligence (Fluid Intelligence/Reasoning Test, n = 28,981; Matrix Pattern Completion, n = 21,038), processing speed (Trail Making Test A, n = 21,257; Symbol Digit Substitution Task, n = 21,056), reaction time (Reaction Time Task, n = 29,318) and working memory (Numeric Memory Test, n = 21,536). All cognitive measures were coded such that higher values indicate better performance. Further details on UK Biobank cognitive measures are described elsewhere (Fawns-Ritchie & Deary, 2020).

#### 2.2.6 Statistical Analyses – UK Biobank

##### 2.2.6.1 Brain structure associations with *FREM3*-GReX

Linear regression models were fitted with *FREM3*-GReX as the independent variable and cortical surface area (SA) as the dependent variable, separately for total SA and regional SA across 62 cortical ROIs. In addition, secondary models were evaluated with a *FREM3*-GReX-by-age interaction. For all models, covariates included sex, age, age^2^, (sex×age), (sex×age^2^), the first 10 genetic principal components, assessment site, and estimated total intracranial volume (eTIV).

Exploratory global and regional analyses with cortical volume (CV) (given its translational relevance to our mouse brain volume measures) and cortical thickness (CT) as dependent variables were also conducted separately, with the same covariates as SA analyses (excluding eTIV for CT). An FDR of 5% was applied across the three global measures and separately across regional comparisons (62 ROIs).

##### 2.2.6.2 Cognitive performance associations with *FREM3*-GReX

For each cognitive measure, separate linear regression models were fitted with *FREM3*-GReX as the independent variable and cognitive performance as the dependent variable. A secondary model was also evaluated with a *FREM3*-GReX-by-age interaction. For both models, covariates included sex, age, age^2^, (sex × age), (sex × age^2^), and the first 10 genetic principal components. An FDR of 5% was applied across the eight cognitive tasks.

Follow-up analyses examined whether total surface area indirectly mediated associations between *FREM3*-GReX and cognitive performance, using the mediate() function from the mediation package in R. For the mediator, linear models were fitted with *FREM3*-GReX as the independent variable and total surface area the dependent variable. Similar outcome models were fitted with cognitive task performance as dependent variables and total surface area as the independent variable. Both models adjusted for the same covariates in the main cognition analyses as well as estimated total intracranial volume (eTIV). Nonparametric bootstrapping using 1,000 samples was used to estimate average causal mediation effects, average direct effects, and total effects. An FDR of 5% was applied across the eight cognitive tasks.

## Supporting information

Supplemental File 1 - Neuroimaging Results

Supplemental File 2 - Human Neuroimaging Results

Supplemental File 3 - Human Cognition Results

## Acknowledgments

YSN was supported by a Koerner New Scientist Award and a Paul Garfinkel New Investigator Catalyst Award administered by the CAMH Foundation, as well as a Discovery Grant from the Natural Sciences and Engineering Research Council of Canada. KPC was supported by the CAMH Koerner Graduate PhD Award.

This research was conducted using the UK Biobank Resource under Application Number 85461.

We thank the CAMH Animal Facility Staff for their support and care of the animals used in this study, and all of the animals that were a part of this study.

## Notes

### Competing Interest Statement

The authors have declared no competing interest.

